## Supplementary figures and movie legend for "Human interictal epileptiform discharges are bidirectional traveling waves echoing ictal discharges"

**Supplemental information:**

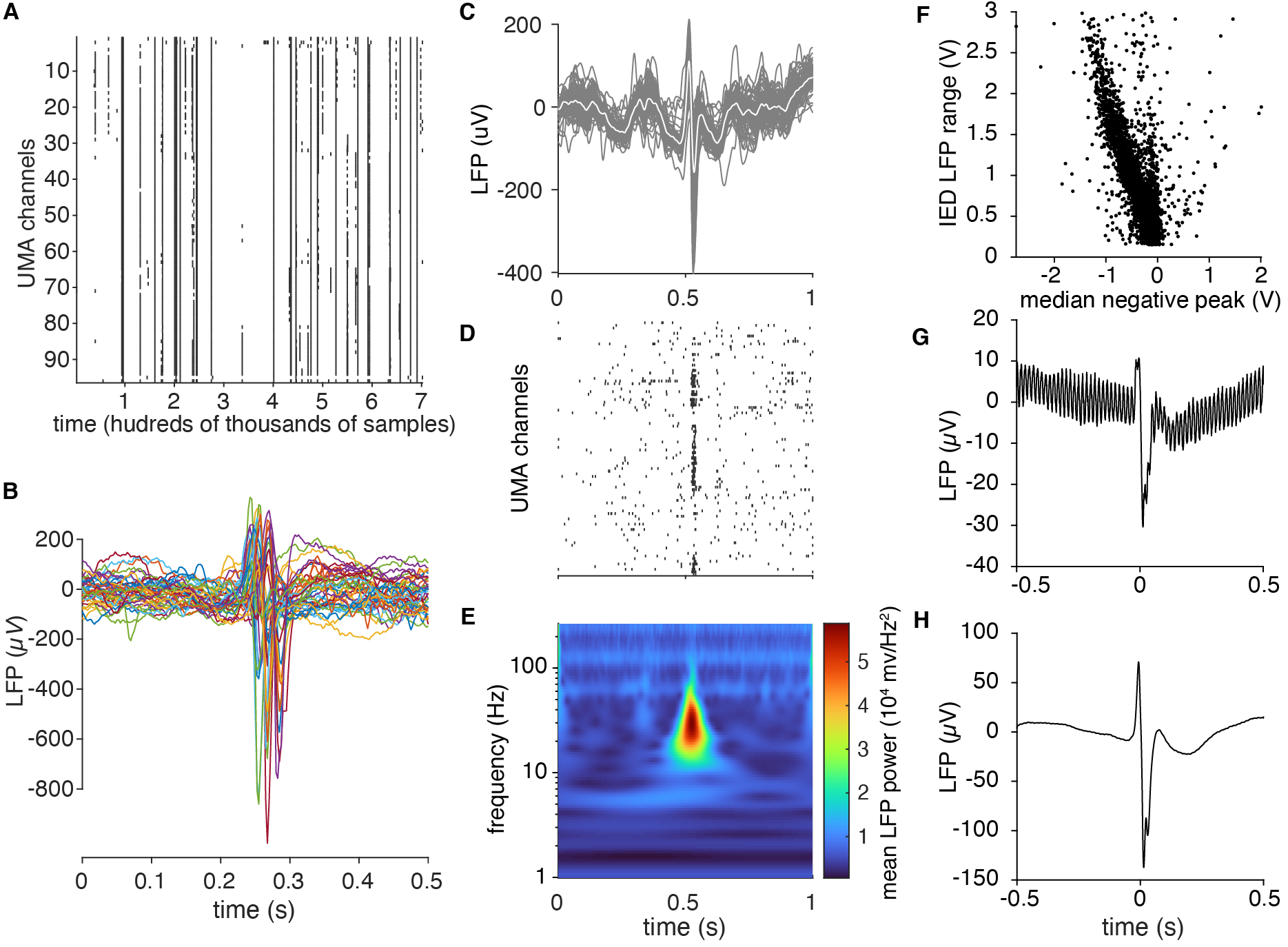
**Figure S1. IED detection and artifact rejection.** (**A**) Raster plot of IED detection times across UEA channels during a segment of data (approximately half an hour duration). (**B**) mean IED waveforms for the retained detections shown in (**A**). These appear as rasters occurring across more than 10 channels in (**A**). (**C**) Recordings of a single IED across all microelectrodes. (**D**) Raster plot of IED-associated MUA firing. (**E**) Mean spectrogram across microelectrodes for the IED shown in (**C**). Prominent Beta power, upon which the detection algorithm operated, is clearly visible. (**F**) A scatter plot of the mean IED negative peak versus the median range of recorded voltages for each IED in one participant after rejecting outliers. (**G**) Mean IED waveform before amplitude rejection. (**H**) Mean IED waveform after IED rejection. (**G**) and (**H**) show that many large amplitude detections were also highly noisy.

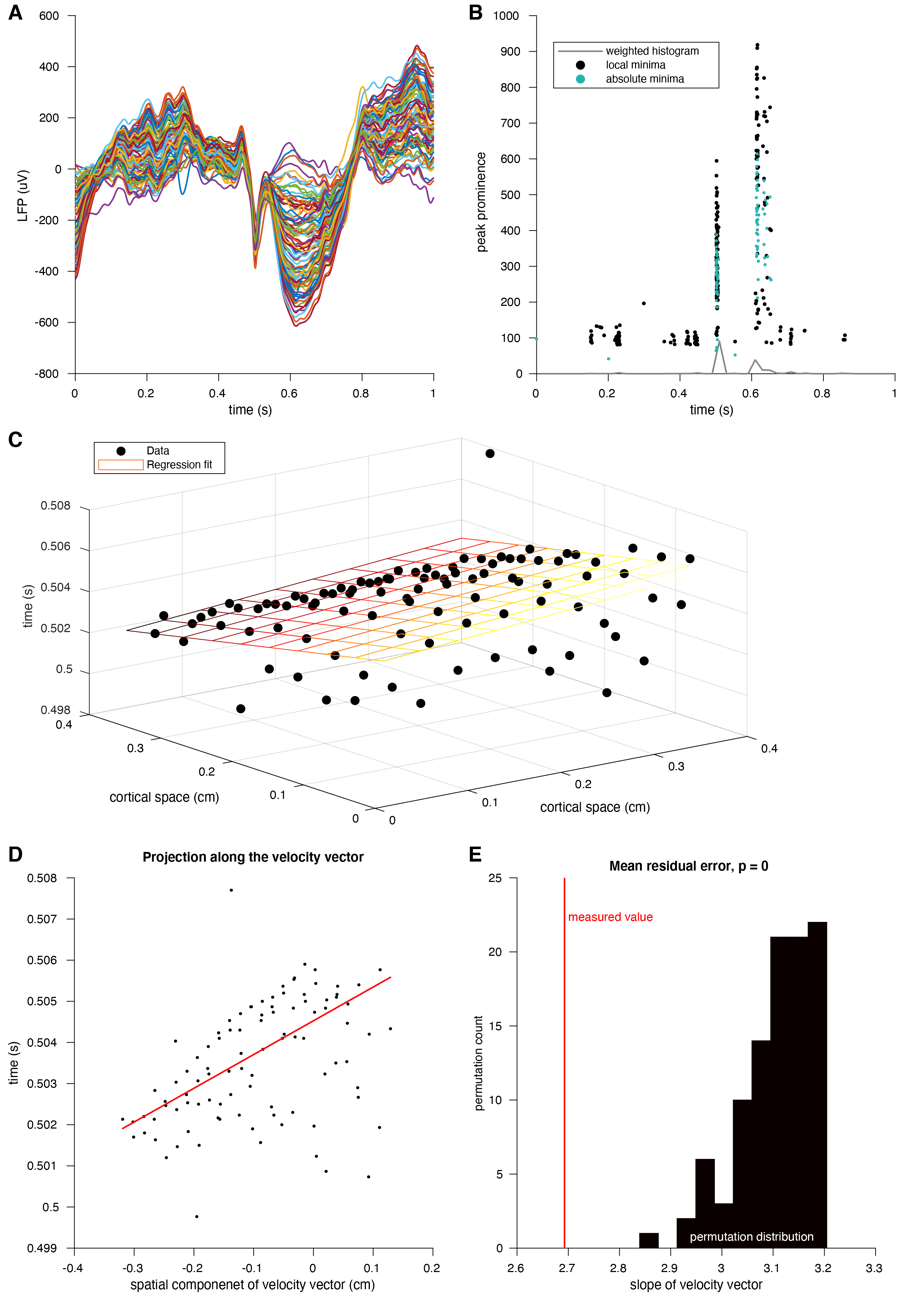

**Figure S2. Classifying IED traveling waves. (A)** Example IED recorded across microelectrodes. Each colored line represents the voltages recorded form a single microelectrode. **(B)** Scatter plot of IED extrema over time. Black dots show local minima and green dots show absolute minima. The weighted histogram of these points that was used to define the time window used for extrema detection is shown in gray. **(C)** Three-dimensional scatter plot of voltage minima timings plotted across the spatial footprint of the UEA. The plane from the multilinear regression model, regularized via least absolute deviation, is shown as a grid in spacetime colored by time of occurrence (earlier: black; intermediate: orange; later: yellow). **(D)** Scatter plot of IED extrema and slope line of the linear regression model projected into two dimensions. **(E)** Visual representation of the permutation test that was used to operationally define IED traveling waves. The true residual absolute deviation is shown in red and residual absolute deviations from spatially shuffled data are shown in black.

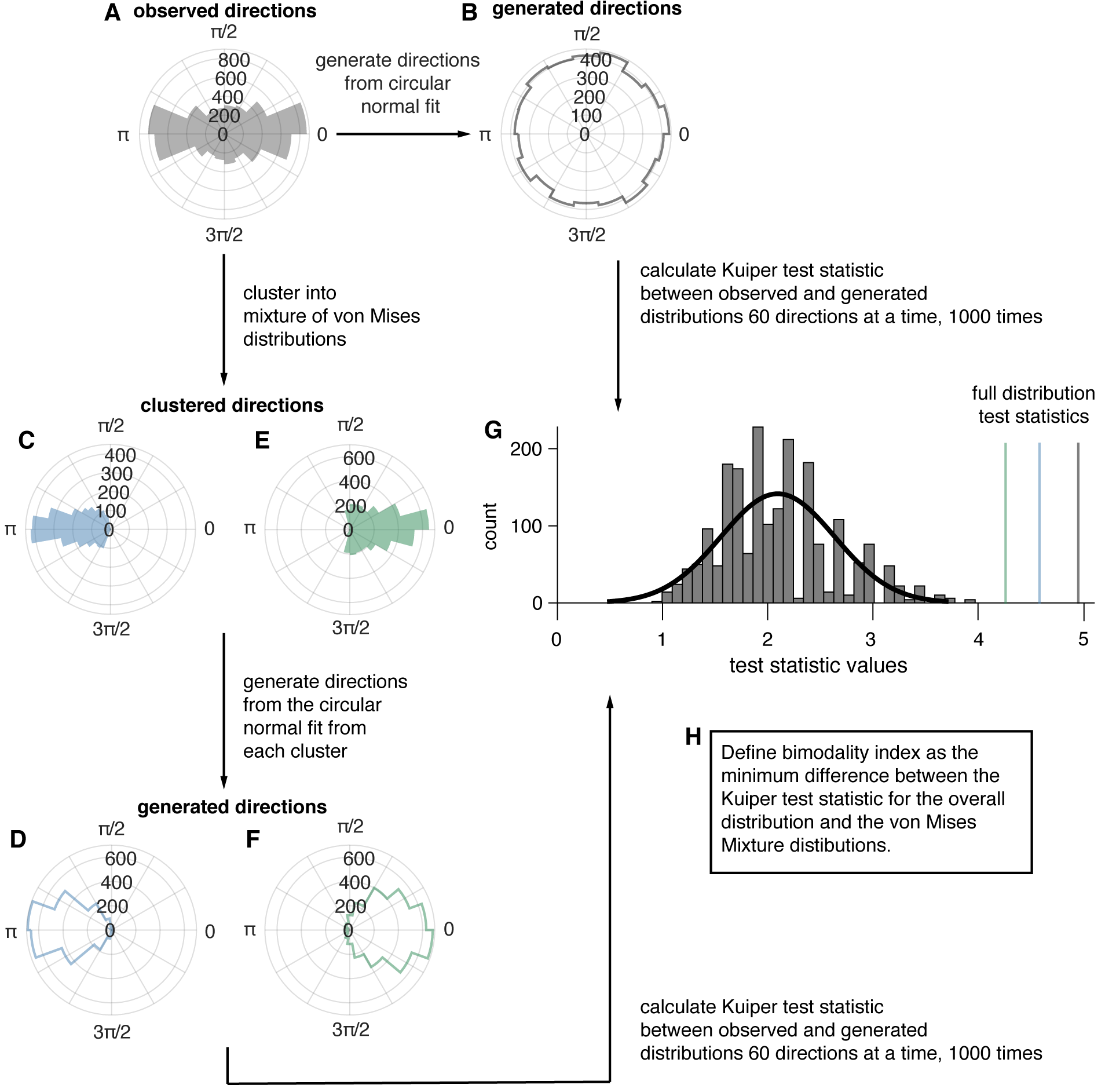

**Figure S3. Procedures for clustering and evaluating the goodness-of-fit of overall and von Mises Mixture distributions.** (**A**) An example IED distribution for one participant. (**B**) a distribution of the same number of directions sampled from the circular normal distribution with the same mean direction, and concentration parameter as the distribution in (**A**). (**C**) The first clustered distribution of angles from (**A**). (**D**) a distribution of the same number of directions sampled from the circular normal distribution with the same mean direction, and concentration parameter as the distribution in (**C**). (**E**) The second clustered distribution of angles from (**A**). (**F**) a distribution of the same number of directions sampled from the circular normal distribution with the same mean direction, and concentration parameter as the distribution in (**C**). (**G**) the permutation distribution of Kuiper test statistics comparing subsamples of angles from real IED distributions and circular normal distributions with the same parameters. The black line shows a gaussian fit to the distribution and colored lines represent the test statistics from the true data distributions. (**H**) A description of the classification rule for unimodal vs bimodal IED distributions.

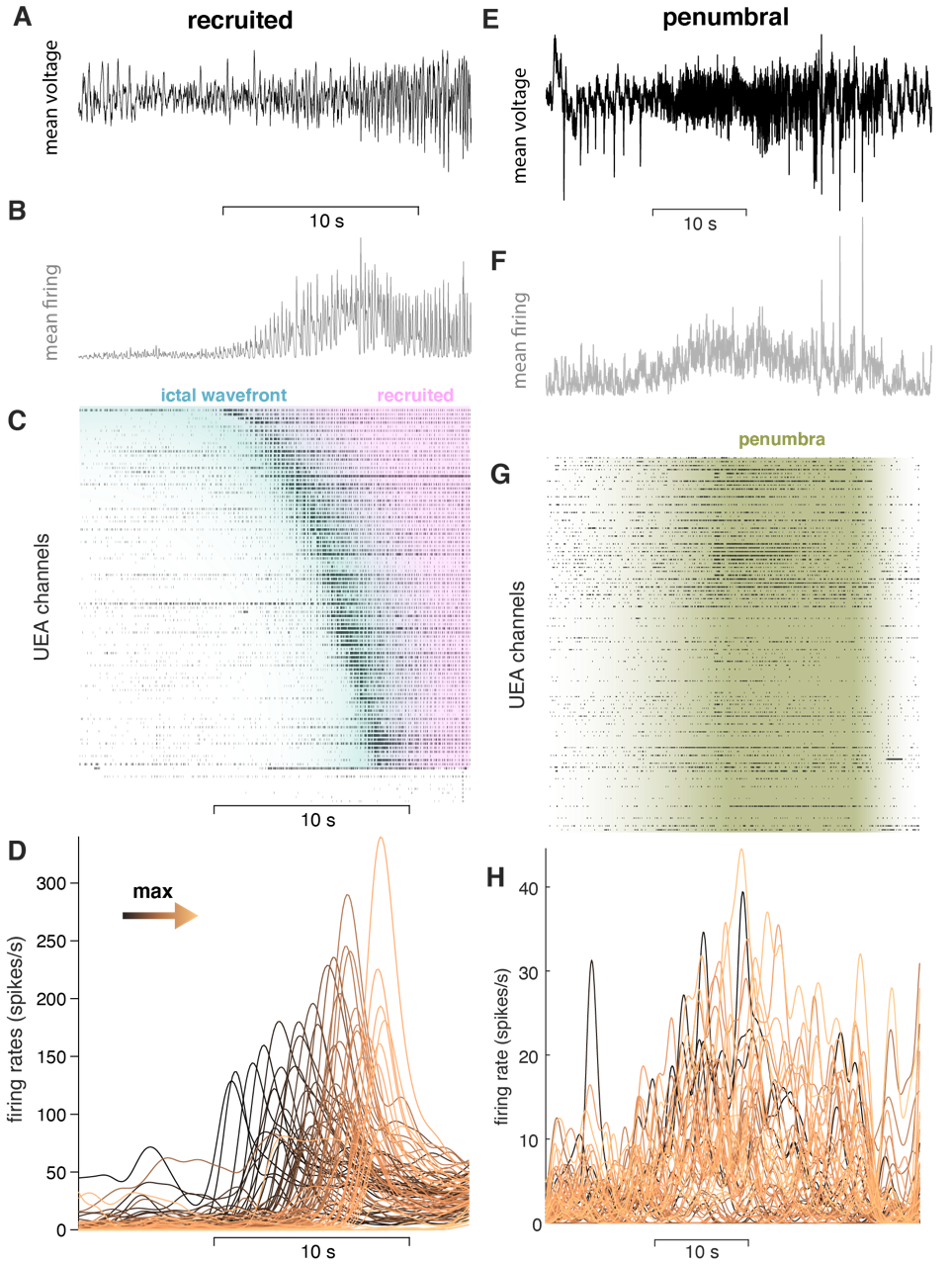

**Figure S4. Examples of each class of microelectrode seizure recording.** (**A**) Mean LFP around the time of UEA recruitment for the “recruited” category. Scale bar shows 10 s. (**B**) Mean MUA firing rate across the microelectrodes for the same time period shown in (**A**). (**C**) Raster plot of MUA event times for each microelectrode on the UEA sorted by recruitment time. (**D**) MUA firing rates on each microelectrode colored by order of recruitment from black to copper. (**E**) Mean LFP for a seizure in the “penumbral” category. Scale bar shows 10 s. (**F**) Mean MUA firing rate across microelectrodes for the same time period shown in (**E**). (**G**) Raster plot of multiunit event times for each microelectrode. (**H**) Multiunit firing rates for each microelectrode colored by the timing of maximal firing during the seizure window. Note the lack of spatiotemporal organization in the “penumbral” example.

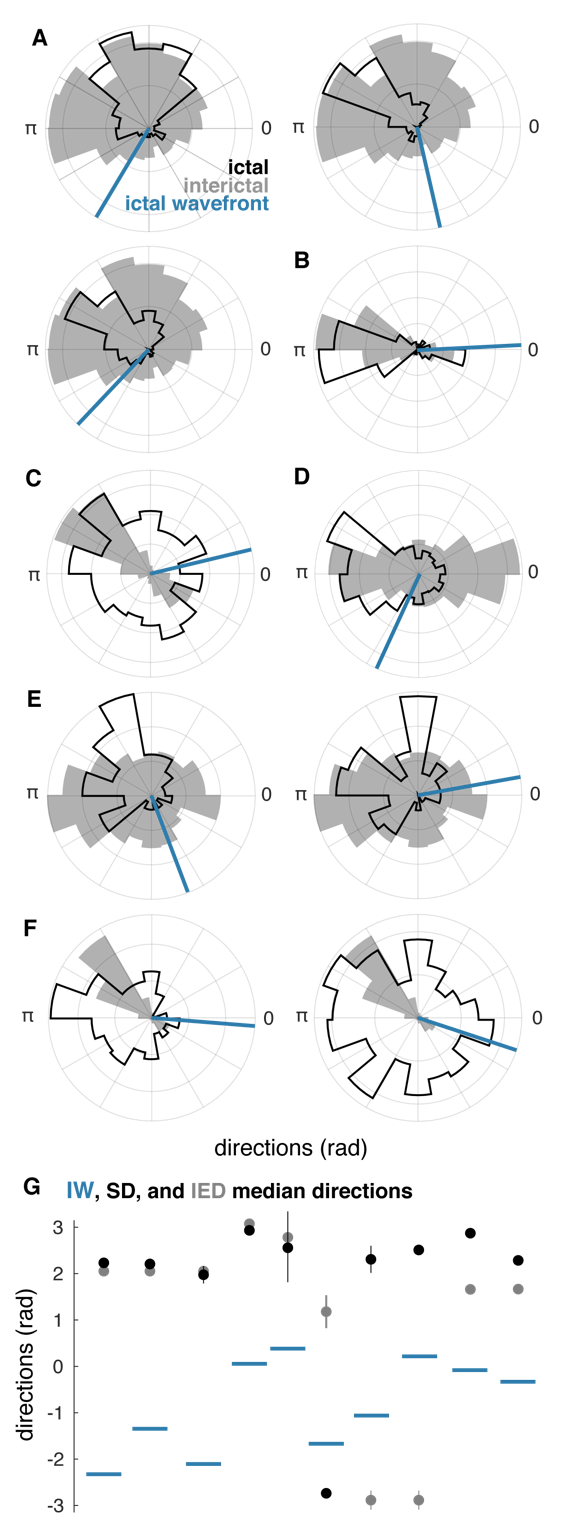

**Figure S5. Raw IED and SD distributions in “recruited” UEA recordings.** (**A-F**) Each lettered subpanel corresponds to one participant. Each polar histogram corresponds to one seizure. Gray histograms show IED distributions. Black histograms show SD distributions, and blue lines show the directions of IWs. (**G**) direction summaries for each seizure ordered and color coded as in (**A-F**). Dots indicate median directions and lines indicate standard deviations.

| **Algorithm S1. Algorithm for detecting IEDs from microelectrode recordings.** | |
| --- | --- |
| **Input:** A matrix of microelectrode voltage recordings, $V(c,t)$, where measured voltage is a function of time, $t$, and which microelectrode array channel it was recorded on, $c$. 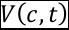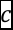 **Output:** A vector of IED times, $I(t)$. 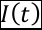 | |
| 1:  2:  3:    4:  5:  6:  7:  8:    9: | **for** each $c$, do: 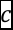 filter data between 20 and 40 Hz (non-causal, 4th order Butterworth filter)  detect peaks, $p$, in filtered signal greater than 8 times the standard deviation of the beta power in the data. 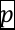 discard peaks that occur within 250 ms of a preceding detection  **for** each $p$, co-occurring within 250 ms, across more than 10 microelectrodes. 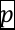 find all local minima of $V(c,t)$ in time, across all $c$. 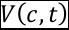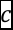 bin local minima in time, across all $c$. 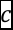 convolve the resulting histogram with a modified Heaviside function,    $H\left( t \right): \left\{ \begin{aligned} 0, 0 \geq n<0.4 \\ 10t, 0.4\geq n<0.5 \\ -t, 0.5\geq n<1 \end{aligned} \right.$ 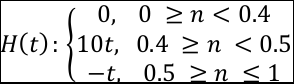 find absolute minima of $V(c,t)$ within the bin containing the most local minima. 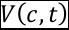 |

**Table S1. Clinical details for research participants.**

| **Participant** | **Age** | **sex** | **Epileptogenic zone** | **UEA implant site** | **Pathology** | **Outcome** |
| --- | --- | --- | --- | --- | --- | --- |
| **1** | 30 | male | right lateral and mesial temporal lobe; nonlesional | Right middle temporal gyrus, 4 cm posterior to the temporal pole | Mesial temporal sclerosis | Engel 2 at 22 months |
| **2** | 25 | male | Left mesial temporal lobe | Left middle temporal gyrus, 3 cm posterior to temporal pole | Non-specific | Engel 1a at 7 months |
| **3** | 32 | male | Right mesial temporal lobe | right inferior frontal gyrus | Normal hippocampus | Engel1a at 2.5 years |
| **4** | 30 | male | left supplementary motor area | left supplementary motor area | N/A (multiple subpial transections performed) | Engel 3 at >2 years |
| **5** | 39 | male | left frontal operculum | left lateral frontal, 2 cm superior to Broca's Area | Nonspecific | Engel 1a at >2 years |
| **6** | 32 | female | left inferior temporal lobe | inferior temporal gyrus, 2.5 cm from temporal pole | 1 neuronal loss; lateral temporal nonspecific | Engel 1a at 55 months |
| **7** | 19 | female | right posteriror lateral temporal | Right posterior temporal, 1cm inferior to angular gyrus | Non-specific | Engel 1a at >2 years |
| **8** | 28 | male | left dorsal posterior prefrontal cortex | left posterior middle frontal gyrus | Mild reactive astrogliosis, patchy microgliosis, Chaslin’s marginal sclerosis | Engel 3a at 32 months |
| **9** | 26 | male | left middle subtemporal | left posterior inferior temporal gyrus | Diffusely infiltrating low grade glioma, IDH-1 negative | Engel 4a at 2 years, 5 months |
| **10** | 30 | male | right middle inferior temporal gyrus | right middle temporal gyrus | Mild astrocytosis | Engel 1a at 12 months |

**Movie S1. Video of IEDs and ictal recruitment.** A movie showing several pre-ictal IEDs propagating in the opposite direction of seizure expansion, in the same direction as SDs. The top trace shows the mean LFP across all microelectrodes for the beginning of one seizure. The lower three panels show three types of data superimposed across the footprint of the microelectrode array. The lower left panel shows LFP voltages recorded on each microelectrode. The lower middle panel shows the slow multiunit firing dynamics, in which the slow propagation of the ictal wavefront can be observed. The lower right panel shows the fast multiunit firing dynamics. The movie progresses at 1/8^th^ the speed of real time and the red bar shows the passage of time, with three seconds indicated by the temporal scale bar.
